# Cognitive Color Coding: Chromatic Tuning Underlying Numerosity Adaptation — Experimental and Factor-Analytic Evidence from Individual Differences

**DOI:** 10.64898/2026.06.27.734875

**Authors:** David Peterzell, Roberto Arrighi, Chiara Di Cesare, Massimo Gurioli, Alessandro Farini, Paolo A. Grasso

**Affiliations:** Department of Physics and Astronomy, University of Florence, Sesto Fiorentino, Italy; Fielding Graduate University, Santa Barbara, CA, USA; Color and Vision Network; Department of Neuroscience, Psychology, Drug Research and Child Health, University of Florence, Florence, Italy; National Research Council, National Institute of Optics, Florence, Italy

**Keywords:** numerosity adaptation, chromatic channels, color discrimination, individual differences, higher-order color mechanisms

## Abstract

Numerosity adaptation (the underestimation of number after exposure to a numerous adaptor) is reduced when adaptor and test differ in color, suggesting that the numerosity system parses items into color-defined categories. Here we ask whether this chromatic selectivity is organized into multiple narrowly tuned chromatic channels, and whether its expression depends on individual chromatic sensitivity. Twenty observers (aged 22–61) completed two psychophysical tasks. First, chromatic discrimination was measured for five hues spaced in 5° CIE L*a*b* steps (ΔH = 0°, 5°, 10°, 15°, 20°) from a red reference (LCh: 54, 118, 38), yielding an individual just-noticeable difference (JND). Second, numerosity adaptation was measured across the same five chromatic distances between a 48-dot adaptor and the test. Observers with superior discrimination (JND < 2.5°) showed robust chromatic tuning, adaptation declining as the test moved away from the adaptor hue, whereas poorer discriminators showed none. Using an interindividual-covariance / factor-analytic approach, we found that adaptation strengths at neighboring chromatic distances were highly correlated and fell off with chromatic separation. Principal component analysis extracted two factors, one loading on the larger chromatic distances and one on the smaller; under oblique (promax) rotation the two factors were substantially correlated (r = .66), implying at least two dissociable but overlapping chromatically tuned mechanisms. These results suggest that numerosity adaptation is mediated by multiple, comparatively narrow chromatic channels, resembling the higher-order color mechanisms inferred from color scaling, SSVEP, and fMRI, rather than the two early cardinal axes (L–M, S–(L+M)).

## 1. Introduction

Humans and many other animals possess a rapid, approximate sense of number that operates without serial counting (Burr & Ross, 2008; Dehaene, 2011). A defining signature of this number sense is its susceptibility to adaptation: after prolonged viewing of a numerous array, a subsequent array appears less numerous, with the opposite effect following adaptation to sparse arrays (Burr & Ross, 2008). Numerosity adaptation has become one of the principal tools for characterizing the mechanisms underlying numerosity perception, and converging psychophysical, EEG, and fMRI evidence locates the adaptable representation at relatively high levels of the visual hierarchy, particularly in parietal cortex, after the stages at which items are grouped and segmented (Castaldi et al., 2016; Cicchini et al., 2016; Grasso et al., 2021a, 2021b, 2022b, 2024; Harvey et al., 2013).

If numerosity is computed after objects have been segmented and bound with their features, then adaptation should be sensitive to the features that define category membership. This prediction has been borne out for color. Grasso et al. (2022) showed that numerosity adaptation is roughly three times stronger when adaptor and test share the same color than when they differ, an effect that holds for physically distinct colors and for physically identical colors rendered perceptually different through a color-assimilation illusion, and that generalizes to pitch in the auditory domain. A subsequent study established that this color selectivity is not an artifact of low-level novelty or perceptual filtering of “old” information, but instead reflects the visual system’s capacity to parse a scene into color-defined categories (Grasso et al., 2025; see also Caponi et al., 2025). Together these findings indicate that numerosity mechanisms have access to color and use it to determine which items belong together. Because the luminance of adaptor and test in these demonstrations was equated only roughly rather than precisely — luminance being a component also found to influence the strength of numerosity adaptation (see Caponi et al., 2025) — the effect should also be established with strictly isoluminant stimuli, so that any contribution of luminance differences to the color-selective adaptation can be excluded.

What remains unknown is the chromatic resolution of this process. Color-selective adaptation has so far been demonstrated only with widely separated colors (e.g., red versus green or blue, separated by roughly 90–135° in color space). Such coarse separations cannot reveal whether the underlying selectivity is carried by a small number of broadly tuned, early cardinal mechanisms (the L–M and S–(L+M) cone-opponent axes) or by multiple, more narrowly tuned mechanisms of the sort increasingly invoked to explain higher-order color appearance. Distinguishing these possibilities requires probing adaptation across small, parametric steps in hue.

A complementary way to characterize sensory mechanisms exploits individual differences. The interindividual-covariance technique, formalized as a “factor-analytic manifesto” by Peterzell (2016; see also Mollon et al., 2017), rests on the premise that performance measures mediated by a common channel should covary across observers more strongly than measures mediated by different channels. Patterns of correlation across stimulus values, summarized by principal component analysis or factor analysis, therefore recover the number and tuning of the underlying mechanisms. The approach has been used to isolate channels for contrast sensitivity, spatial frequency, motion, stereopsis, and color (Peterzell, 2016; Peterzell et al., 2017; Serrano-Pedraza et al., 2018). In the chromatic domain it has been especially productive: factor analyses of hue scaling indicate that color appearance depends on multiple narrowly tuned mechanisms that vary independently of one another and are inconsistent with a simple two-axis opponent code (Emery, Volbrecht, Peterzell, & Webster, 2017a, 2017b, 2023). Convergent evidence from steady-state visual evoked potentials points to broadly distributed, intermediate hue tuning rather than strictly cardinal mechanisms (Kaneko, Kuriki, Andersen, & Peterzell, 2025; Rozman et al., 2025; Chen & Gegenfurtner, 2021), and a long tradition of work on unique-hue variability reaches similar conclusions (Bosten, 2022; Bosten & Boehm, 2014).

Critically for the present study, the same covariance logic has recently been brought to bear on number itself. Anobile et al. (2024) measured the precision with which observers reproduced symbolic numbers as sequences of taps, correlated precision across target numbers between observers, and found correlations that were highest for neighboring numbers and fell off with numerical distance. Factor analysis and PCA on these data returned two bell-shaped components tuned to low and high numbers, providing the first individual-differences evidence for numerosity-selective channels. That study establishes both the feasibility and the interpretive framework for applying covariance analysis to numerosity, but it concerned the number-to-action transformation and made no contact with color. The same covariance technique has since been carried beyond the number-to-action transformation to purely sensory visual numerosity: Petrizzo et al. (2025) recovered tuned channels for spatial (dot-array) and temporal (flash-sequence) numerosity, and for sensorimotor number, finding shared tuning between the temporal visual and sensorimotor tasks but at most partly shared mechanisms for spatial numerosity. Unlike the present work, however, these individual-differences studies of number have not examined chromatic structure.

Here we combine these strands. We measured numerosity adaptation across five small chromatic steps from a fixed red reference, in observers whose chromatic discrimination thresholds for exactly those colors were independently determined. This design allows three questions to be addressed. First, does chromatic tuning of numerosity adaptation survive at fine chromatic scales? Second, does the expression of that tuning depend on an observer’s chromatic sensitivity, such that only good discriminators reveal it? Third, and most novel, when the covariance of adaptation strength across chromatic distance is analyzed across observers, how many tuned chromatic mechanisms are implied, and what is their tuning? On the basis of the higher-order color literature we predicted that adaptation would be carried by more than the two cardinal mechanisms, and that interindividual covariance would reveal at least two dissociable chromatic factors.

## 2. Methods

### 2.1. Participants

Twenty adults (9 female, 11 male; aged 22–61 years, mean 33.6, SD 14.1) with normal or corrected-to-normal acuity and normal color vision took part. Testing was carried out at the Department of Physics and Astronomy, University of Florence (Sesto Fiorentino). All participants gave informed consent and procedures conformed to the Declaration of Helsinki.

### 2.2. Apparatus and stimuli

Stimuli were generated in MATLAB with the Psychophysics Toolbox and presented on a monitor (1366 × 768) in a dimly lit room, viewed binocularly at approximately 57 cm. All stimuli were clouds of non-overlapping dots drawn within a circular region of 8° diameter and presented for 200 ms at 10° eccentricity from a central fixation cross, with dot positions randomized on every presentation.

Five chromaticities were defined in CIE L*a*b* space, beginning from a reference red (LCh: 54, 118, 38) and rotating in 5° hue steps to yield colors at chromatic distances of ΔH = 0°, 5°, 10°, 15°, and 20° from the reference (Figure 1). The five colors varied only in hue: they were equiluminant, at approximately 38 cd/m^2^ (measured with a Minolta LS-110 photometer), with saturation held constant. Because the set spans only a 20° arc of hue, the four non-reference colors are perceptually close to the reference red, becoming progressively more orange with increasing ΔH.

**Figure 1.**
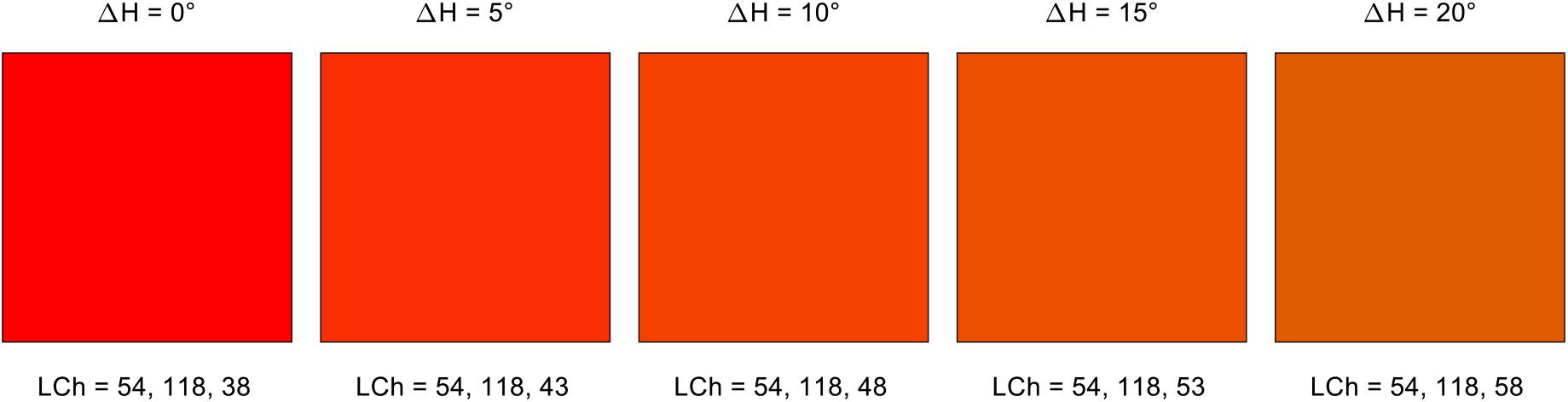
The five equiluminant stimulus colors, shown as patches with their chromatic distance (ΔH) from the red reference (LCh: 54, 118, 38) and their LCh values. The colors vary only in hue; saturation (C) and luminance (L) were held constant.

On each trial the dots forming a stimulus were a single color drawn from this set. The reference color throughout was the red at ΔH = 0° (LCh: 54, 118, 38); the other colors served as comparison or test colors at the four non-zero chromatic distances.

### 2.3. Procedure

The experiment comprised two tasks, illustrated in Figure 2. In both, observers fixated a central cross and dot clouds appeared briefly (200 ms) in the periphery; observers responded with key presses and received no trial-by-trial feedback.

**Figure 2.**
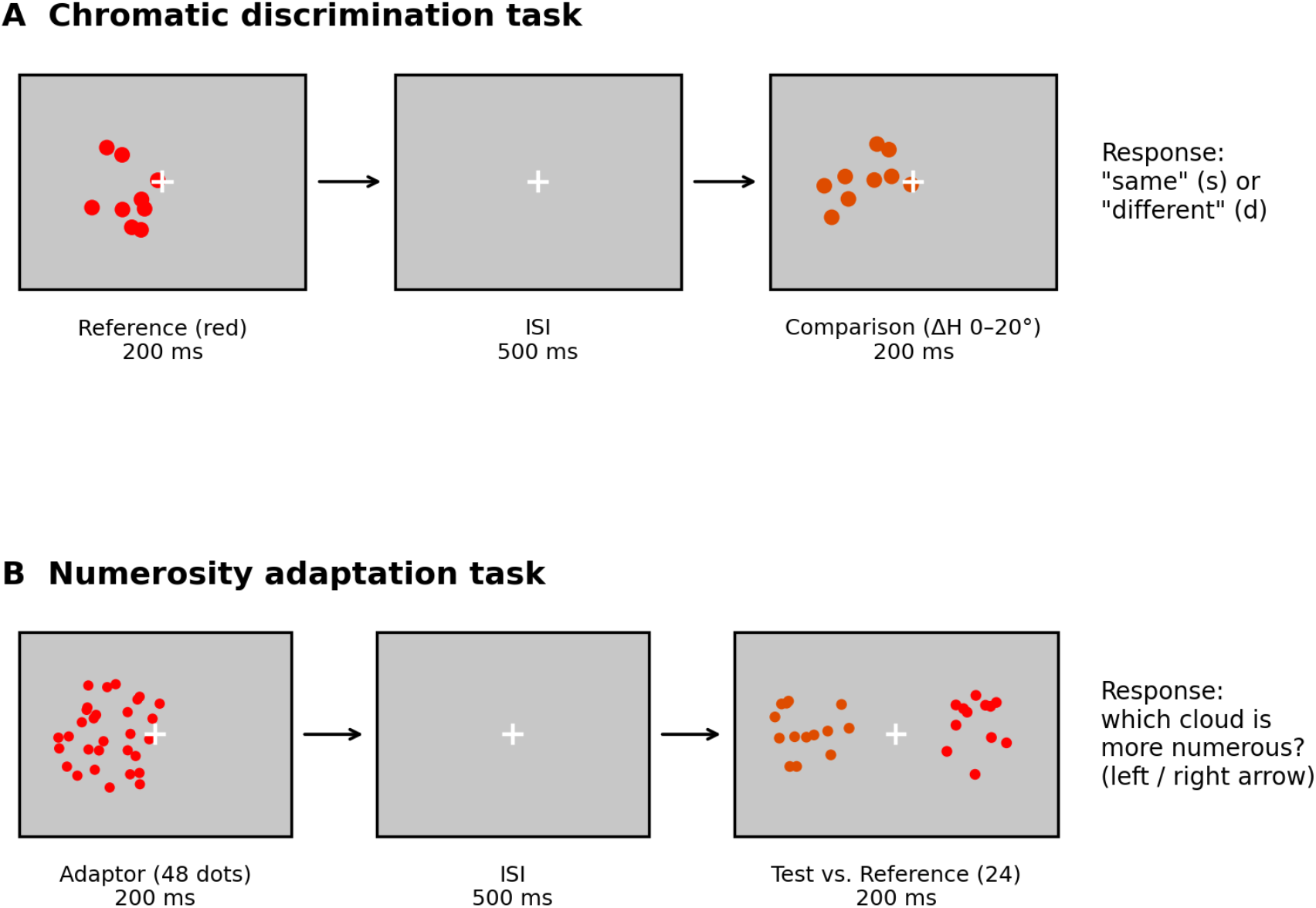
Schematic of the two tasks and how observers interacted with the stimuli. (A) Chromatic discrimination: a red reference cloud was followed, after a 500-ms interval, by a second cloud that was either identical in hue or displaced by one of the four chromatic steps; observers reported “same” or “different.” (B) Numerosity adaptation: a 48-dot adaptor was followed by the simultaneous presentation of a variable-numerosity test and a 24-dot reference; observers reported which cloud was more numerous. Adaptor–test chromatic distance was varied across the same five levels. Timings and stimulus sizes are illustrative.

Task 1 (chromatic discrimination). Two dot clouds were presented sequentially on the left of the display. The first was always the reference red (LCh: 54, 118, 38); the second was either chromatically identical to the reference or differed from it by one of the four hue steps. Observers reported whether the second cloud was the same (“s”) or different (“d”) from the first. Each of the five conditions was presented 20 times (100 trials total). The proportion of “different” responses was plotted against chromatic distance and fit with a cumulative function; the just-noticeable difference (JND; calculated as half the difference between the 75% and 25% thresholds) was taken as the chromatic distance supporting threshold discrimination, providing a per-observer index of chromatic sensitivity.

Task 2 (numerosity adaptation across chromatic distance). Adaptation was measured with a two-alternative comparison procedure. A trial began with a 48-dot adaptor on the left (1000 ms), followed after a 500-ms inter-stimulus interval by the simultaneous presentation (200 ms) of a test and a reference cloud, left and right of fixation. The reference always contained 24 dots; the test varied across 12, 14, 17, 20, 24, 29, 34, 40, and 48 dots. Observers indicated which cloud was more numerous. The chromatic distance between adaptor and test was manipulated across the same five levels used in Task 1 (ΔH = 0–20°). A baseline was obtained with the identical procedure but without the adaptor. For each observer and chromatic condition, the proportion of “test more numerous” responses was fit with a cumulative Gaussian to estimate the point of subjective equality (PSE); adaptation magnitude was computed as the PSE shift from baseline, normalized by the reference numerosity and expressed as a percentage.

### 2.4. Individual-differences analysis

For the covariance analysis, each observer contributed five adaptation-magnitude values (one per chromatic distance), which were standardized (z-scored) across observers within each condition. The 20 × 5 matrix was submitted to a Pearson correlation analysis, and the resulting 5 × 5 correlation matrix was examined for the near-diagonal structure expected if neighboring chromatic distances share tuned mechanisms. Dimensionality was assessed with a scree plot of the eigenvalues, and the structure was characterized with principal component analysis followed by varimax (orthogonal) rotation, retaining two components. Observers were additionally classified as good or poor chromatic discriminators for grouped analyses. For each observer, the JND obtained from the chromatic-discrimination psychometric function in Task 1 served as an index of chromatic sensitivity, with lower JNDs indicating finer discrimination. Observers whose JND exceeded 2.5° were classified as poor discriminators and the remainder as good discriminators; this criterion yielded 15 good and 5 poor discriminators.

## 3. Results

### 3.1. Mean effects

Averaged across observers, chromatic discrimination improved steeply with chromatic distance: observers reported a difference on fewer than 10% of trials when the two clouds were identical (ΔH 0°), on roughly 30% at ΔH 5°, and on about 90% by ΔH 10°, reaching ceiling at ΔH 15–20°. On the basis of these JND measurements, observers were classified as good (JND < 2.5°; n = 15) or poor (JND > 2.5°; n = 5) chromatic discriminators, a distinction we draw on throughout the analyses that follow.

Adaptation produced a robust overall underestimation of test numerosity: PSEs during adaptation were consistently higher than baseline across all chromatic conditions. The critical question was whether adaptation magnitude varied with chromatic distance between adaptor and test, and whether this depended on chromatic sensitivity. Figure 3 shows mean adaptation magnitude as a function of chromatic distance, separately for good and poor discriminators. For good discriminators (Figure 3, left), adaptation was largest when adaptor and test shared the same hue and declined monotonically as chromatic distance increased, describing a chromatic tuning gradient. For poor discriminators (Figure 3, right), adaptation showed no systematic ordering by ΔH, varying irregularly across conditions. The same dissociation was evident in the psychometric functions: clearly separated and ΔH-ordered for good discriminators, but superimposed for poor discriminators.

**Figure 3.**
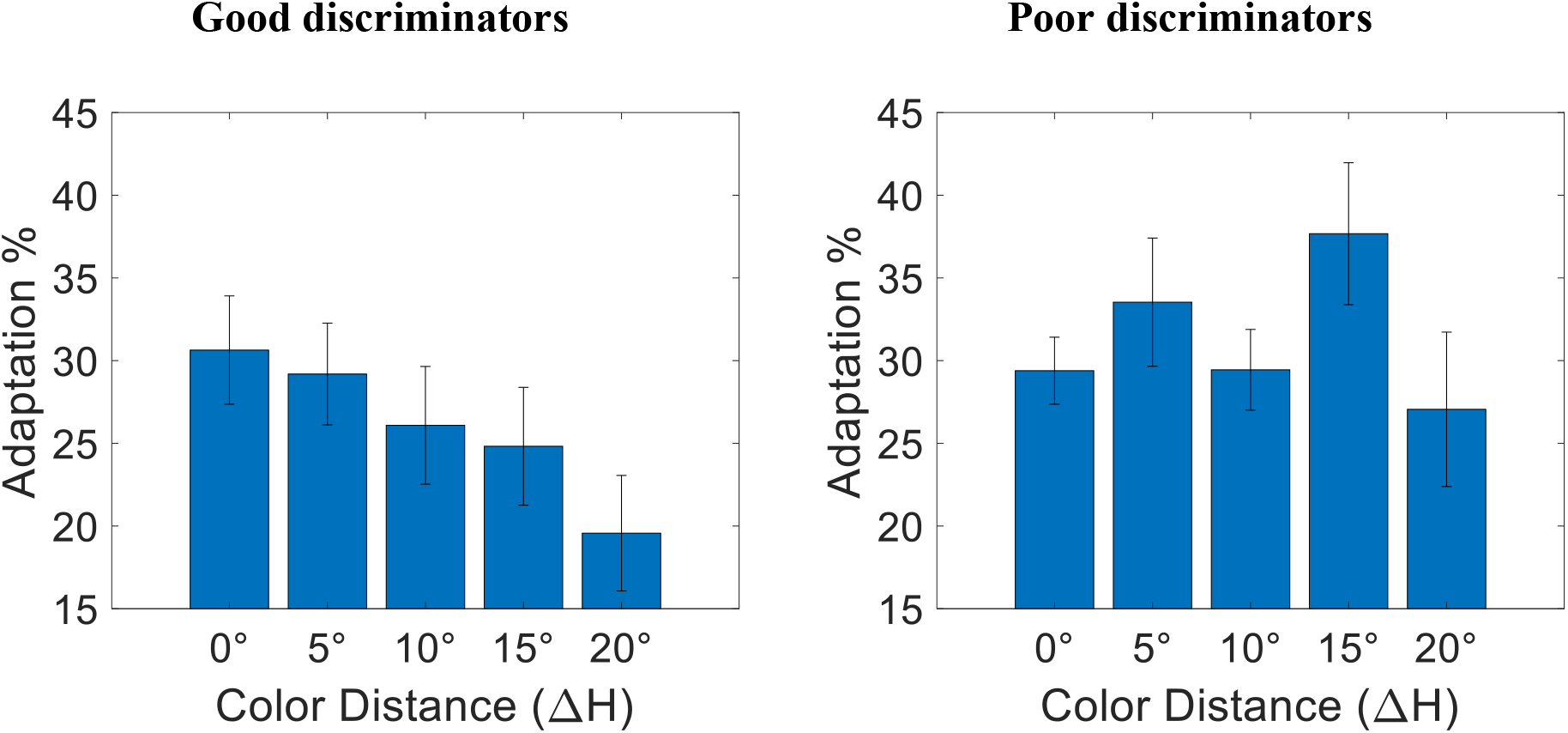
Mean numerosity-adaptation magnitude (percentage PSE shift from baseline) as a function of adaptor–test chromatic distance, for good discriminators (left) and poor discriminators (right).

### 3.2. Individual differences: correlations and factor analysis

We next applied the interindividual-covariance approach to the standardized adaptation magnitudes. The correlation matrix (Table 1) showed the structure expected if adaptation at neighboring chromatic distances is mediated by shared tuned mechanisms: correlations were highest for adjacent chromatic distances (ΔH 0–5°: r = .83; ΔH 10–15°: r = .83) and decreased with chromatic separation, reaching r = .54 for the most widely separated pair (ΔH 0–20°). This near-diagonal, distance-dependent fall-off is the qualitative signature of multiple overlapping channels rather than a single common mechanism, which would predict an unstructured matrix (Anobile et al., 2024; Peterzell, 2016).

**Table 1.**
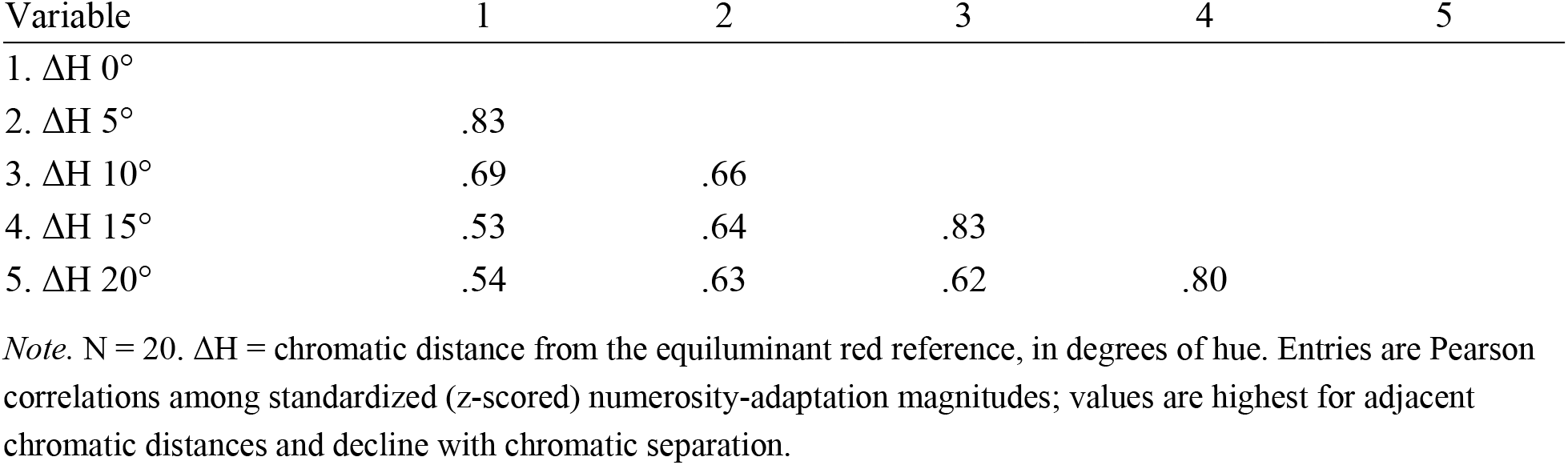
Correlations Among Standardized Numerosity-Adaptation Magnitudes Across the Five Chromatic Distances.

A scree plot of the eigenvalues (Figure 4) showed a single dominant component (eigenvalue 3.71) followed by an elbow. A straight scree line, running from an eigenvalue of 0.67 at the first component to zero at the fifth, passes below the second component (0.67) yet tracks the third and subsequent values (0.38, 0.17, 0.08); by this scree criterion two components are retained. This is reinforced by the systematic near-diagonal structure of the correlation matrix and mirrors the interpretive logic of comparable covariance studies, in which PCA recovers the minimal number of components consistent with the data rather than an exhaustive count of channels (Anobile et al., 2024). The first two components together accounted for 87.5% of the total variance (74.1% and 13.3%, respectively), and the communalities were uniformly high after extraction (.79 to .94).

**Figure 4.**
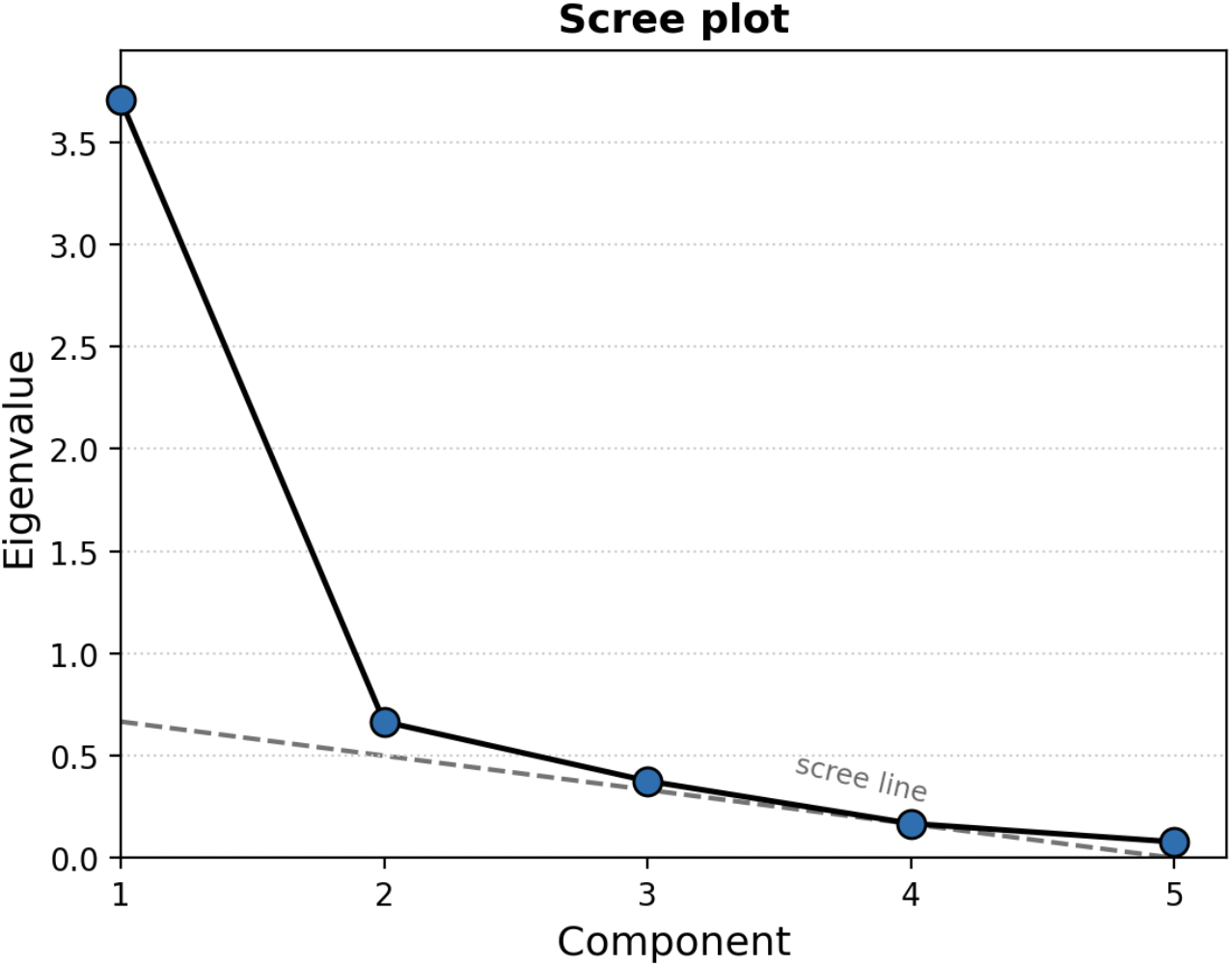
Scree plot of the eigenvalues from the principal component analysis of adaptation magnitudes across the five chromatic distances. A single dominant component is followed by an elbow. The dashed scree line runs from an eigenvalue of 0.67 at the first component (on the vertical axis) to zero at the fifth component (on the horizontal axis); the second component falls just above this line whereas the third and later components lie on it, supporting retention of two components for rotation.

Principal component analysis with varimax (orthogonal) rotation yielded two interpretable factors (Figure 5); these are principal component analyses with rotation rather than common-factor analyses. After rotation the two factors accounted for 46.8% and 40.7% of the variance. Factor 1 loaded most strongly on the larger chromatic distances (.92 at ΔH 15° and .85 at ΔH 20°, with ΔH 10° intermediate at .72), whereas Factor 2 loaded most strongly on the smaller chromatic distances (.93 at ΔH 0° and .85 at ΔH 5°). The two factors crossed over near ΔH 10°, which loaded moderately on both, producing the complementary, oppositely sloped loading profiles characteristic of two overlapping tuned mechanisms. This pattern indicates dissociable processing of subtle versus larger chromatic variations and is consistent with the existence of at least two dissociable, comparatively narrowly tuned chromatic mechanisms underlying color-selective numerosity adaptation.

**Figure 5.**
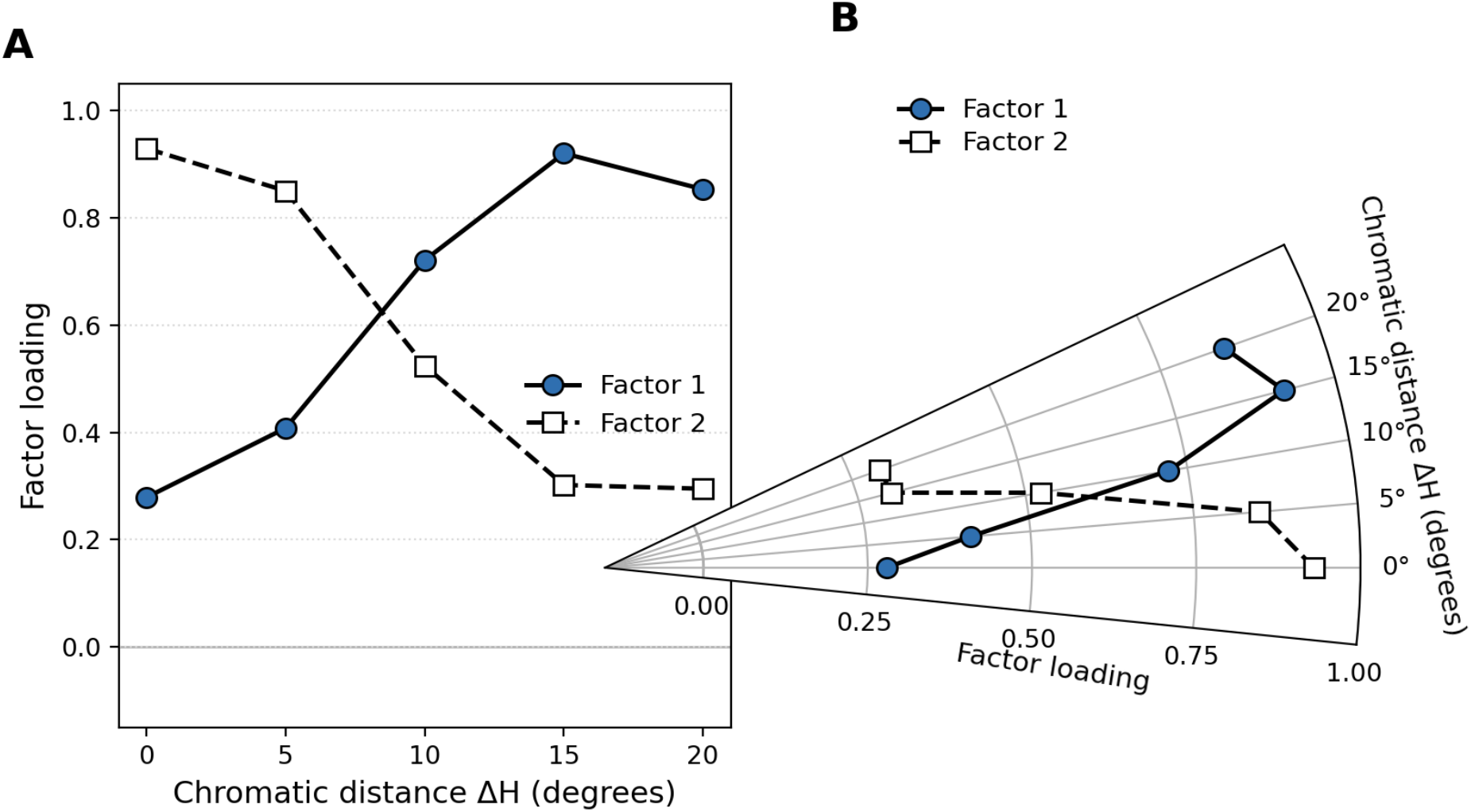
Varimax-rotated loadings of the two factors as a function of chromatic distance (ΔH), shown in (A) Cartesian and (B) polar coordinates. In the polar panel the radial axis, from the origin outward, is the factor loading, and the angular position is the chromatic distance ΔH. Factor 1 (filled symbols) increases with chromatic distance, loading on ΔH 10–20°; Factor 2 (open symbols) decreases, loading on ΔH 0–5°. The two profiles cross near ΔH 10°, the signature of two overlapping, oppositely tuned chromatic mechanisms. The loading axis extends slightly below zero so that the coordinates match those of Figure 6.

Because two finely spaced chromatic mechanisms are unlikely to be strictly independent, we repeated the analysis with promax (oblique) rotation, which permits the factors to correlate (Figure 6). The oblique solution reproduced the structure of the varimax solution: Factor 1 again loaded on the larger chromatic distances and Factor 2 on the smaller distances, with ΔH 10° loading primarily on Factor 1 and the two profiles crossing near ΔH 10°. Critically, the two factors were themselves substantially correlated (r = .66), indicating that they are not independent but interdependent, as would be expected of overlapping chromatic channels with adjacent tuning. Two pattern loadings slightly exceeded 1.0 and were truncated to 1.0 for plotting.

**Figure 6.**
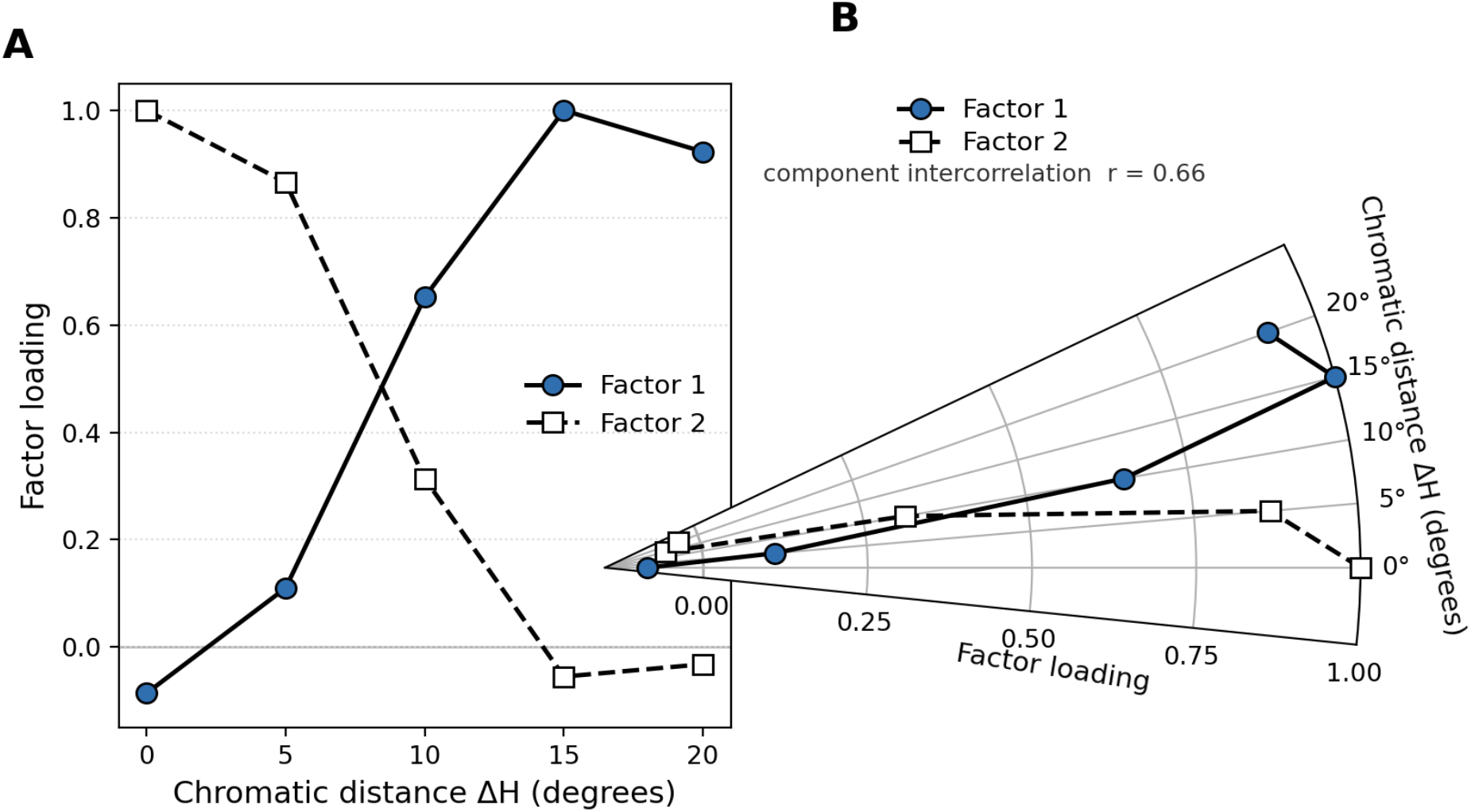
Loadings from the principal component analysis with promax (oblique) rotation, shown in (A) Cartesian and (B) polar coordinates on the same axes as Figure 5 (radial axis, from the origin outward, = factor loading; angular position = chromatic distance ΔH). Factor 1 (filled symbols) loads on the larger chromatic distances and Factor 2 (open symbols) on the smaller distances, reproducing the crossover seen under varimax rotation. Pattern loadings above 1.0 were truncated to 1.0 for display, and the slightly negative loadings (near ΔH 0° for Factor 1 and ΔH 15–20° for Factor 2) fall just inside the zero line. The two components were correlated (r = .66, the component intercorrelation from the oblique rotation), confirming that the factors are not independent.

## 4. Discussion

We measured numerosity adaptation across five small chromatic steps from a fixed red reference and related it, observer by observer, to chromatic discrimination thresholds for the same colors. Three results emerged. First, chromatic tuning of numerosity adaptation persists at fine chromatic scales: among observers who could discriminate the small hue steps, adaptation declined systematically as the test hue moved away from the adaptor. Second, this tuning depended on chromatic sensitivity: it was clear in good discriminators and absent in poor discriminators, whose adaptation profiles were flat and whose psychometric functions overlapped. Third, and most novel, interindividual covariance in adaptation strength across chromatic distance revealed near-diagonal correlation structure and two factors, one tuned to smaller and one to larger chromatic distances; under oblique rotation these factors were themselves correlated, implying at least two dissociable but overlapping chromatically tuned mechanisms.

That numerosity adaptation is tuned to color at all constrains where, in the visual hierarchy, numerosity is computed. A representation that can be rendered color-selective must have access to the features that bind items into sets, which places the adaptable numerosity signal after figures have been segmented from the background and grouped by their surface properties, rather than at the earliest, feature-blind stages of vision. Consistent with this, color-selective adaptation tracks the perceived rather than the physical color of the adaptor (Grasso et al., 2022a), is associated with a late-component modulation in VEP (Grasso et al., 2022b), acts on perceived rather than physical numerosity (Fornaciai et al., 2016), and numerosity is represented in topographic maps in parietal cortex (Harvey et al., 2013). On this view the chromatic tuning reported here reflects a relatively high-level, salient-feature parsing of the scene into color-defined subsets, of the kind that would let the visual system enumerate items of one kind among distractors of another (Caponi et al., 2025; Grasso et al., 2025).

The dependence of tuning on chromatic sensitivity is informative rather than merely a power difference. If color-selective adaptation is carried by mechanisms with access to fine chromatic differences, then observers who cannot resolve those differences should fail to show selectivity, precisely the dissociation we observed. Poor discriminators still adapted (their overall PSEs shifted), but their adaptation was not organized by hue, as expected if the categorization signal that normally modulates adaptation is unavailable when the relevant chromatic differences are below threshold. This pattern reinforces the interpretation that color-selective numerosity adaptation reflects genuine chromatic categorization (Grasso et al., 2022a, 2025) rather than a low-level by-product of local stimulus change (cf. Yousif et al., 2024).

The individual-differences analysis is the central contribution. Following the covariance logic that has isolated channels for contrast, spatial frequency, motion, stereopsis, and color (Peterzell, 2016; Mollon et al., 2017; Reynaud & Hess, 2017), and that Anobile et al. (2024) recently extended to number, we treated each observer’s profile of adaptation across chromatic distance as a set of measures whose covariance reveals shared mechanisms. The highest correlations fell on neighboring chromatic distances and declined with separation, the qualitative pattern predicted by multiple overlapping channels and not by a single mechanism, and PCA returned two tuned factors. As in the sensorimotor-number case, the two-factor solution should be read as evidence for at least two dissociable mechanisms rather than as an exact count; PCA summarizes covariance with the minimal number of components, and finer channels could be smoothed into a coarser two-factor description (Anobile et al., 2024, 2026; Petrizzo et al., 2025).

Although the structure of chromatic tuning, rather than the reality of numerosity adaptation, is our focus, the present results bear on a continuing debate about what numerosity aftereffects measure. On one account the aftereffect reflects the discounting of locally repeated, “old” information rather than abstract number (Yousif et al., 2024). Several features of the color-selective effect are difficult to reconcile with these alternatives. Selectivity follows perceived rather than physical color (Grasso et al., 2022a); it survives a control experiment in which positional local novelties were strictly controlled (Grasso et al., 2025); and it is graded by stimulus identity defined by low-level features rather than by any salient change: a highly visible change of motion does not reduce adaptation, whereas a change of color does (Caponi et al., 2025; Burr et al., 2025). At the same time, a non-selective component of adaptation persists when adaptor and test colors differ, on the order of 10 to 15% in published reports (Grasso et al., 2022a, 2025), suggesting that part of the phenomenon might reflect the contribution of a more general mechanism extracting numerical information regardless of stimulus identity. The dependence of chromatic tuning on numerosity adaptation in our view suggests a genuine chromatic categorization of the stimuli.

The number and form of these factors bear on where color-selective numerosity processing sits. Two early cardinal mechanisms, the L–M and S–(L+M) cone-opponent axes, do not naturally predict graded tuning across a series of small hue steps confined to one region of color space; a cardinal account would more readily predict an abrupt change once a cardinal boundary is crossed. The graded fall-off and the two overlapping factors we observe align better with the multiple, comparatively narrowly tuned higher-order chromatic mechanisms inferred from hue-scaling individual differences (Emery et al., 2017a, 2017b, 2023), from SSVEP measures of cortical color tuning (Kaneko et al., 2025; Rozman et al., 2025; Chen & Gegenfurtner, 2021), and from unique-hue variability and other non-cardinal dimensions of color appearance, such as the warm–cool axis (Bosten, 2022; Bosten & Boehm, 2014; Manalansan, Whitehead, & Webster, 2025). This progression has a direct parallel in the spatial domain, where spatial-frequency and orientation channels become more sharply tuned as one ascends the visual hierarchy: the broadly tuned, magnocellular- and parvocellular-like mechanisms of the retina and lateral geniculate nucleus give way to the far more narrowly tuned spatial-frequency and orientation channels of striate cortex (Derrington & Lennie, 1984; De Valois, Albrecht, & Thorell, 1982; Hubel & Wiesel, 1968; Wilson, McFarlane, & Phillips, 1983). On this view, the color information that gates numerosity adaptation is read out from a population of narrowly tuned chromatic units of the kind associated with higher-order color appearance and parietal numerosity processing, rather than from the two early cardinal channels.

Several limitations qualify these conclusions. The sample is modest for factor analysis (n = 20), and the grouped good/poor comparisons should be treated as suggestive; replication with a larger and more sensitivity-balanced sample is warranted. The colors sampled only a narrow arc of color space near red, which was deliberate (the aim was to probe fine chromatic resolution), but it leaves open whether comparable tuning, and a comparable number of factors, would be recovered around other reference hues or across the full hue circle. The two-factor solution rests partly on the scree criterion, and we have been explicit that it indexes a lower bound on the number of mechanisms. Equally, these data cannot settle the bandwidth of the chromatic mechanisms involved: hue-scaling studies favor narrowly tuned higher-order mechanisms (Emery et al., 2017a, 2017b, 2023), whereas steady-state evoked-potential work points to more broadly tuned, intermediate mechanisms (Chen & Gegenfurtner, 2021; Rozman et al., 2025; but see Kaneko et al., 2025), and an adaptation-based covariance result is consistent with multiple beyond-cardinal mechanisms without fixing their tuning width. Because the sampled hues span only a narrow red-to-orange arc, the two factors also cannot be firmly attributed to distinct higher-order mechanisms as opposed to finer structure within the cardinal red–green system; sampling the full hue circle will be needed to distinguish these accounts. Finally, the covariance technique recovers mechanisms shared across observers and is insensitive to idiosyncratic, observer-specific channels (Mollon et al., 2017).

Notwithstanding these limits, the results extend color-selective numerosity adaptation from coarse to fine chromatic scales, show that its expression is gated by individual chromatic sensitivity, and, by importing the interindividual-covariance framework into this domain, provide the first individual-differences evidence consistent with numerosity adaptation drawing on more than one comparatively narrow, higher-order chromatic mechanism. They thereby connect the literatures on numerosity adaptation, higher-order color mechanisms, and the factor-analytic study of sensory channels, and suggest that the visual system’s coding of “how many” is intertwined, at a relatively high level, with a finely resolved code for “what color.”

## Supporting information

Data used in study

## Acknowledgements

We are grateful to Chiara Di Cesare, our co-author whose undergraduate thesis (Di Cesare, 2024) provided data and some analyses that contributed to the present study, including the chromatic-discrimination measurements and the classification of observers into good and poor discriminators. D.H.P. was partially supported in 2025 as a Visiting Professor in the Department of Physics and Astronomy, University of Florence.

## Data availability

The data are available in the Harvard Dataverse at https://doi.org/10.7910/DVN/EVMLHV.

## Use of AI-assisted technologies in manuscript preparation

During the preparation of this work, the authors used Claude (Anthropic) to assist with literature mapping, reference formatting, and refinement of wording for clarity. All original research, analyses, arguments, and interpretations were developed by the authors, who reviewed and edited all AI-assisted text and take full responsibility for the content of this publication.

## References

Anobile, G., Martelli, S., Bieber, E., Tinelli, F., & Ditano, S. (2026). Sensorimotor numerosity uniquely supports arithmetic development in children. i-Perception, 17(3), 1–14. 10.1177/20416695261443209

Anobile, G., Petrizzo, I., Paiardini, D., Burr, D. C., & Cicchini, G. M. (2024). Sensorimotor mechanisms selective to numerosity derived from individual differences. eLife, 12, RP92169. 10.7554/eLife.92169

Bosten, J. M. (2022). Do you see what I see? Diversity in human color perception. Annual Review of Vision Science, 8(1), 101–133. 10.1146/annurev-vision-093020-112820

Bosten, J. M., & Boehm, A. E. (2014). Empirical evidence for unique hues? Journal of the Optical Society of America A, 31(4), A385–A393. 10.1364/JOSAA.31.00A385

Burr, D., Anobile, G., & Arrighi, R. (2025). Number adaptation: Reply. Cognition, 254, 105870. 10.1016/j.cognition.2024.105870

Burr, D., & Ross, J. (2008). A visual sense of number. Current Biology, 18(6), 425–428. 10.1016/j.cub.2008.02.052

Caponi, C., Castaldi, E., Grasso, P. A., & Arrighi, R. (2025). Feature-selective adaptation of numerosity perception. Proceedings of the Royal Society B, 292(2039), 20241841. 10.1098/rspb.2024.1841

Castaldi, E., Aagten-Murphy, D., Tosetti, M., Burr, D., & Morrone, M. C. (2016). Effects of adaptation on numerosity decoding in the human brain. NeuroImage, 143, 364–377. 10.1016/j.neuroimage.2016.09.020

Chen, J., & Gegenfurtner, K. R. (2021). Electrophysiological evidence for higher-level chromatic mechanisms in humans. Journal of Vision, 21(8), 12. 10.1167/jov.21.8.12

Cicchini, G. M., Anobile, G., & Burr, D. C. (2016). Spontaneous perception of numerosity in humans. Nature Communications, 7, 12536. 10.1038/ncomms12536

Dehaene, S. (2011). The number sense: How the mind creates mathematics (Rev. ed.). Oxford University Press.

Derrington, A. M., & Lennie, P. (1984). Spatial and temporal contrast sensitivities of neurones in lateral geniculate nucleus of macaque. The Journal of Physiology, 357, 219–240. 10.1113/jphysiol.1984.sp015498

De Valois, R. L., Albrecht, D. G., & Thorell, L. G. (1982). Spatial frequency selectivity of cells in macaque visual cortex. Vision Research, 22(5), 545–559. 10.1016/0042-6989(82)90113-4

Di Cesare, C. (2024). Influenza della sensibilità cromatica su fenomeni di adattamento percettivo [Undergraduate thesis]. University of Florence.

Emery, K. J., Volbrecht, V. J., Peterzell, D. H., & Webster, M. A. (2017a). Variations in normal color vision. VI. Factors underlying individual differences in hue scaling and their implications for models of color appearance. Vision Research, 141, 51–65. 10.1016/j.visres.2016.12.006

Emery, K. J., Volbrecht, V. J., Peterzell, D. H., & Webster, M. A. (2017b). Variations in normal color vision. VII. Relationships between color naming and hue scaling. Vision Research, 141, 66–75. 10.1016/j.visres.2016.12.007

Emery, K. J., Volbrecht, V. J., Peterzell, D. H., & Webster, M. A. (2023). Fundamentally different representations of color and motion revealed by individual differences in perceptual scaling. Proceedings of the National Academy of Sciences USA, 120(4), e2202262120. 10.1073/pnas.2202262120

Fornaciai, M., Cicchini, G. M., & Burr, D. C. (2016). Adaptation to number operates on perceived rather than physical numerosity. Cognition, 151, 63–67. 10.1016/j.cognition.2016.03.006

Grasso, P. A., Anobile, G., Arrighi, R., Burr, D. C., & Cicchini, G. M. (2022a). Numerosity perception is tuned to salient environmental features. iScience, 25(4), 104104. 10.1016/j.isci.2022.104104

Grasso, P. A., Anobile, G., & Arrighi, R. (2021a). Numerosity adaptation partly depends on the allocation of implicit numerosity-contingent visuo-spatial attention. Journal of Vision, 21(1), 12. 10.1167/jov.21.1.12

Grasso, P. A., Anobile, G., Caponi, C., & Arrighi, R. (2021b). Implicit visuospatial attention shapes numerosity adaptation and perception. Journal of Vision, 21(8), 26. 10.1167/jov.21.8.26

Grasso, P. A., Petrizzo, I., Caponi, C., Anobile, G., & Arrighi, R. (2022b). Visual P2p component responds to perceived numerosity. Frontiers in Human Neuroscience, 16, 1014703. 10.3389/fnhum.2022.1014703

Grasso, P. A., Petrizzo, I., Coniglio, F., & Arrighi, R. (2024). Electrophysiological correlates of temporal numerosity adaptation. Frontiers in Neuroscience, 18, 1349540. 10.3389/fnins.2024.1349540

Grasso, P. A., Anobile, G., Gurioli, M., Cicchini, G. M., & Arrighi, R. (2025). Color-selective numerosity adaptation depends on the automatic categorization of colored information. iScience, 28(6), 112572. 10.1016/j.isci.2025.112572

Harvey, B. M., Klein, B. P., Petridou, N., & Dumoulin, S. O. (2013). Topographic representation of numerosity in the human parietal cortex. Science, 341, 1123–1126. 10.1126/science.1239052

Hubel, D. H., & Wiesel, T. N. (1968). Receptive fields and functional architecture of monkey striate cortex. The Journal of Physiology, 195(1), 215–243. 10.1113/jphysiol.1968.sp008455

Kaneko, S., Kuriki, I., Andersen, S. K., & Peterzell, D. H. (2025). Individual variability in steady-state VEP responses for hues sweeping around cardinal color axes: Clues to cortical color coding? Journal of Vision, 25(12), 2. 10.1167/jov.25.12.2

Manalansan, J., Whitehead, L. A., & Webster, M. A. (2025). Warm versus cool colors and their relation to color perception. Journal of Vision, 25(4), 13. 10.1167/jov.25.4.13

Mollon, J. D., Bosten, J. M., Peterzell, D. H., & Webster, M. A. (2017). Individual differences in visual science: What can be learned and what is good experimental practice? Vision Research, 141, 4–15. 10.1016/j.visres.2017.11.001

Peterzell, D. H. (2016). Discovering sensory processes using individual differences: A review and factor-analytic manifesto. Electronic Imaging, 2016(16), 1–11. 10.2352/ISSN.2470-1173.2016.16.HVEI-112

Peterzell, D. H. (2026). Supplementary: Data for “Cognitive color coding: Chromatic tuning underlying numerosity adaptation — Experimental and factor-analytic evidence from individual differences” (Peterzell et al., 2026) [Data set]. Harvard Dataverse. 10.7910/DVN/EVMLHV

Peterzell, D. H., Serrano-Pedraza, I., Widdall, M., & Read, J. C. A. (2017). Thresholds for sine-wave corrugations defined by binocular disparity in random dot stereograms: Factor analysis of individual differences reveals two stereoscopic mechanisms tuned for spatial frequency. Vision Research, 141, 127–135. 10.1016/j.visres.2017.11.002

Petrizzo, I., Cicchini, G. M., Burr, D. C., & Anobile, G. (2025). Multisensory number channels derived from individual differences. Multisensory Research, 38(6–8), 383–402. 10.1163/22134808-bja10154

Reynaud, A., & Hess, R. F. (2017). Characterization of spatial frequency channels underlying disparity sensitivity by factor analysis of population data. Frontiers in Computational Neuroscience, 11, 63. 10.3389/fncom.2017.00063

Rozman, A., Watts, D. J., Somers, L. P., Günel, B., Racey, C., Barnes, K., & Bosten, J. M. (2025). Tuning of cortical color mechanisms revealed using steady-state visually evoked potentials. Imaging Neuroscience, 3, IMAG.a.130. 10.1162/IMAG.a.130

Serrano-Pedraza, I., Boegaerts, D., Read, J. C. A., & Peterzell, D. H. (2018). Orientation tuning for spatial vision and stereopsis: Factor analysis of individual differences in contrast and disparity thresholds. Journal of Vision, 18(10), 126. 10.1167/18.10.126

Wilson, H. R., McFarlane, D. K., & Phillips, G. C. (1983). Spatial frequency tuning of orientation selective units estimated by oblique masking. Vision Research, 23(9), 873–882. 10.1016/0042-6989(83)90055-X

Yousif, S. R., Clarke, S., & Brannon, E. M. (2024). Number adaptation: A critical look. Cognition, 249, 105813. 10.1016/j.cognition.2024.105813

