## Supplementary material for "Cognitive Color Coding: Chromatic Tuning Underlying Numerosity Adaptation — Experimental and Factor-Analytic Evidence from Individual Differences": Data used in study

Peterzell, D. H. (2026). *Supplementary: Data for "Cognitive color coding: Chromatic tuning underlying numerosity adaptation — Experimental and factor-analytic evidence from individual differences" (Peterzell et al., 2026)* [Data set]. Harvard Dataverse. https://doi.org/10.7910/DVN/EVMLHV

| | A | B | C | D | E | F |
|---|---|---|---|---|---|---|
| 1 | **deltaH (0)** | **deltaH (5)** | **deltaH (10)** | **deltaH (15)** | **deltaH (20)** | **JND** |
| 2 | 1.379 | 1.515 | 1.209 | -0.077 | -1.231 | 0.820 |
| 3 | 0.365 | -0.771 | -1.447 | -1.782 | -1.264 | 0.972 |
| 4 | 0.199 | 0.616 | 0.056 | 1.556 | 1.250 | 1.173 |
| 5 | -0.312 | -0.230 | -0.789 | -0.575 | -0.646 | 0.795 |
| 6 | -0.186 | 0.206 | 0.541 | 1.060 | 0.696 | 1.169 |
| 7 | 0.664 | 0.344 | -0.037 | -0.404 | 0.681 | 0.424 |
| 8 | -0.601 | -0.562 | -1.199 | -0.497 | -0.262 | 0.542 |
| 9 | -0.778 | -1.346 | -0.151 | -0.354 | -0.927 | 0.691 |
| 10 | 0.467 | 1.193 | 0.802 | 0.830 | 1.053 | 1.432 |
| 11 | -1.046 | -0.918 | -1.754 | -1.892 | -0.686 | 0.942 |
| 12 | 0.498 | 0.113 | -0.147 | -0.184 | -0.073 | 0.033 |
| 13 | -0.489 | -0.928 | -0.257 | -0.207 | -0.352 | 1.023 |
| 14 | -0.553 | 0.260 | 0.045 | 0.147 | 0.731 | 0.990 |
| 15 | 1.818 | 1.877 | 1.296 | 1.602 | 1.234 | 0.971 |
| 16 | 0.978 | 0.203 | 0.461 | 0.143 | -0.121 | 0.033 |
| 17 | -2.475 | -1.792 | -1.405 | -1.580 | -2.168 | 0.722 |
| 18 | 0.225 | -0.755 | 1.914 | 1.184 | 0.615 | 0.033 |
| 19 | -1.018 | -0.819 | -0.300 | 0.091 | 0.263 | 0.581 |
| 20 | -0.413 | 0.374 | -0.108 | 0.237 | -0.486 | 2.003 |
| 21 | 1.278 | 1.423 | 1.273 | 0.701 | 1.693 | 0.665 |
